# Astrocytic Ceruloplasmin Deficiency Triggers Iron Toxicity and Neurodegeneration in a LRRK2 Parkinson’s Tri-Culture Model

**DOI:** 10.1101/2025.08.01.668099

**Authors:** Veronica Testa, Jara Montero-Muñoz, Valentina Baruffi, Marieke Alzeer, Loris Mularoni, Irene Fernández-Carasa, Giulia Simmini, Núria Comellas-Comaposada, Franz Arnold Ake, Yvonne Richaud, Christin Weissleder, Josep Amengual, Isabel Fabregat, Michela Deleidi, Mireya Plass, Stefano Pluchino, Angel Raya, Antonella Consiglio

## Abstract

Astrocytes and microglia carrying the LRRK2-G2019S mutation contribute to non-cell-autonomous dopaminergic neuron (DAn) degeneration in Parkinson’s disease (PD), but the mechanisms underlying their interplay remain unclear. Here, we developed a novel induced pluripotent stem cell (iPSC)-derived tri-culture system comprising healthy DAn and either LRRK2-mutant or isogenic control iPSC-derived astrocytes and microglia. Using integrated functional assays and transcriptomic profiling, we found that mutant astrocytes adopt a hyperreactive state, driving microglial activation and subsequent DAn degeneration. Mechanistically, we identified a selective downregulation of ceruloplasmin (CP), a copper-dependent ferroxidase, in mutant astrocytes, leading to disrupted iron homeostasis with accumulation of Fe^2+^ and ROS. This iron dysregulation mediated both microglial reactivity and neurodegeneration. Notably, pharmacological restoration of CP re-established iron homeostasis, reduced microglial activation, and protected DAn from degeneration. Our findings uncover a novel astrocyte-microglia-neuron axis driving PD pathogenesis and showcase the power of our unique stem cell tri-culture platform for dissecting disease mechanisms and discovering therapeutic targets.

## Introduction

Parkinson’s Disease (PD), the most prevalent neurodegenerative movement disorder, affects over 6.1 million individuals worldwide^1,2^. The disease is characterized by the progressive loss of dopaminergic neurons (DAn) in the *substantia nigra pars compacta* (SNpc), and the presence of intraneuronal protein aggregates known as Lewy Bodies^3–5^. PD is increasingly recognized as a multifactorial disorder, driven by a complex interplay of pathogenic mechanisms including neuroinflammation, mitochondrial dysfunction, oxidative stress, and protein aggregation. Notably, brain iron accumulation, particularly in the SNpc, is a consistent neuropathological feature of PD^6–9^. Iron dysregulation is thought to exacerbate neurodegeneration by promoting oxidative damage via Fenton chemistry, generating cytotoxic free radicals^7^. Despite advances in understanding PD pathogenesis, disease-modifying therapies remain elusive, underscoring the critical need to identify novel molecular mechanisms and therapeutic targets.

While most PD cases are sporadic, approximately 15% are linked to genetic mutations, with *LRRK2* variants being the most common^10^. Given the clinical and pathological overlap between *LRRK2*-linked and sporadic PD^11^, studying *LRRK2* mutations provides a powerful tool for elucidating broadly relevant disease mechanisms. *LRRK2* encodes a large, ubiquitously expressed protein with both GTPase and protein kinase domains. The G2019S mutation, located in the protein kinase domain, hyperactivates LRRK2 and disrupts cellular homeostasis^12,13^.

Historically, PD research focused on DAn dysfunction, but glial cells are now recognized as key contributors to pathogenesis. Reactive astrocytes are consistently observed in post-mortem PD brains^14^, and hiPSC-based models of *LRRK2*-PD astrocytes recapitulate disease phenotypes, including α-synuclein accumulation, metabolic deficits, impaired chaperone-mediated autophagy (CMA), increased cytokine secretion, and morphological alterations^15,16^. These astrocytes exacerbate blood-brain barrier dysfunction and DAn loss in co-culture systems^15,16^, demonstrating their non-cell-autonomous toxicity.

Similarly, microglia (MG) isolated from PD patients exhibit a pro-inflammatory phenotype in the SNpc^17–19^. In *LRRK2*-PD models, mutant MG show hypermotility, excessive phagocytosis, and elevated ROS production, alongside lysosomal and autophagic dysfunction^20–23^, particularly after pro-inflammatory stimulation. These altered phenotypes contribute to DAn degeneration, further implicating MG in PD progression. Taken together, these findings underscore the importance of astrocyte-MG interactions in PD, but existing models fail to capture their reciprocal crosstalk in a human-relevant system. Addressing this gap is critical for uncovering new therapeutic avenues.

To bridge this gap, we developed a novel tri-culture system combining hiPSC-derived astrocytes, MG, and DAn, where astrocytes and MG harbour the *LRRK2*-G2019S mutation (or isogenic controls), and DAn are from healthy individuals (CTRL). Using this platform in conjunction with single-cell RNA sequencing (scRNA-seq), we identified *LRRK2*-PD astrocytes as key drivers of MG activation and DAn degeneration. Transcriptional profiling revealed a selective downregulation of ceruloplasmin (CP) in mutant astrocytes, leading to iron accumulation and ROS generation that propagate neurotoxicity. Critically, restoring CP levels pharmacologically re-established iron homeostasis, suppressed MG reactivity, and protected DAn from neurodegeneration. These findings define a new astrocyte-MG-DAn axis in LRRK2-PD and nominate CP as a therapeutic target.

## Results

### A tri-culture system for modelling neuro-glia interactions in Parkinson’s Disease

To investigate the impact of glial cell crosstalk on DAn survival in PD, we developed a tri-culture system comprising astrocytes, MG, and DAn (**Figure 1A**). Astrocytes and MG were differentiated from two independent *LRRK2*-PD iPSC lines (*LRRK2*-PD1 and *LRRK2*-PD2) and their isogenic controls (ISO-PD1 and ISO-PD2) using established protocols^20,24^. Glial identity was confirmed by assessing the expression of GFAP in mature astrocytes and IBA1 in MG (**Figure S1A**). DAn were derived from a healthy control iPSC line (CTRL) using previously described protocols^25–27^. By day 18, DAn expressed the classical ventral midbrain (vm) DAn fate markers LMX1A and FOXA2. At day 35 of differentiation, DAn were considered mature, as they expressed TH and FOXA2, along with the mature neuronal marker MAP2. The expression of TH and MAP2 within DAn was maintained until day 70, the latest timepoint in our experimental setup (**Figure S1B, Table 1**).

**Figure 1.**
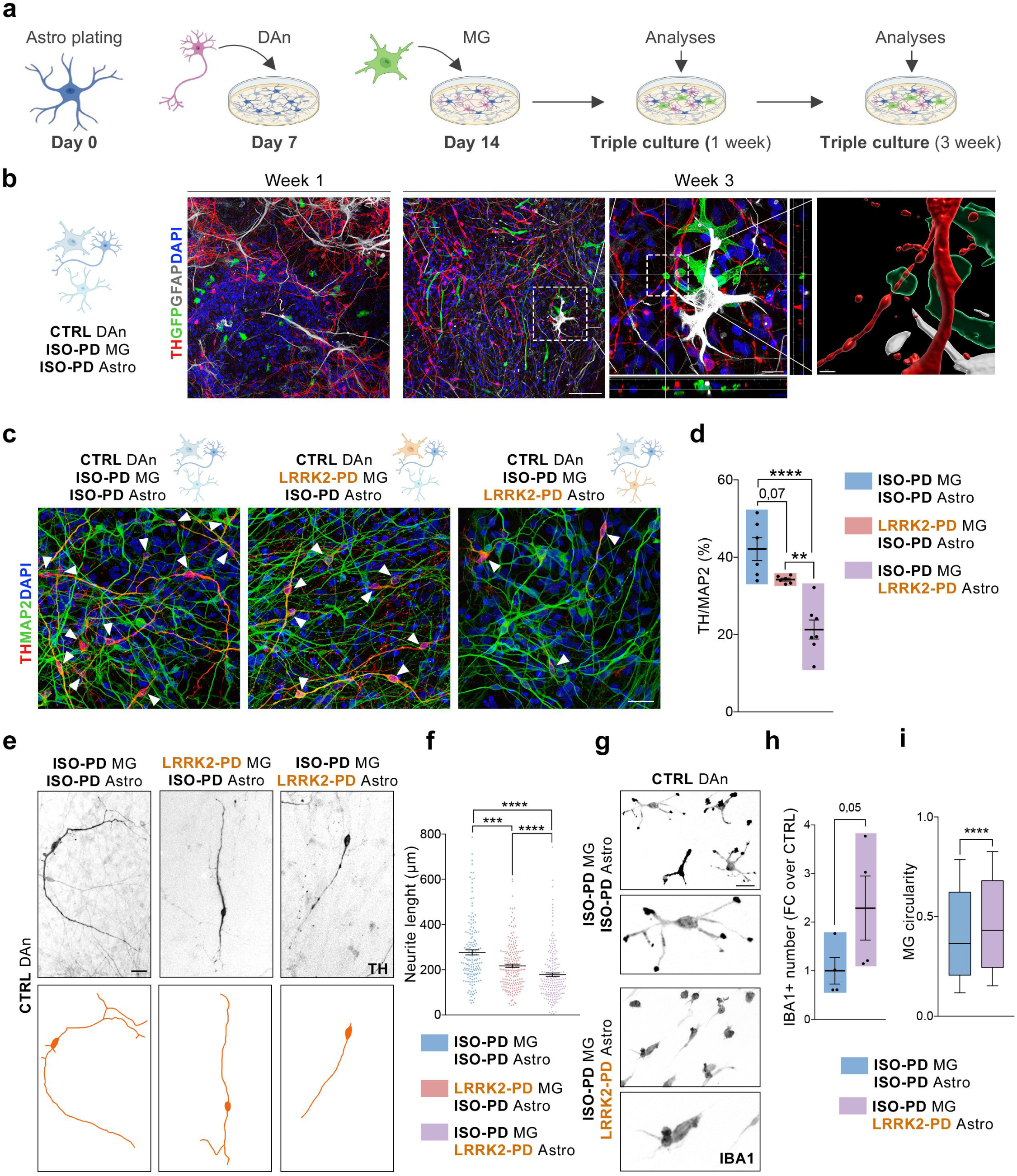
Generation and characterisation of the tri-culture system. **a.** Schematic representation of tri-culture generation and timepoints for analyses. **b.** Representative ICC for tri-culture after 1 or 3 weeks, displaying the presence of TH+ DAn (CTRL: SP11), GFP+ MG (ISO-PD2: SP12 wt/wt), and GFAP+ astrocytes (ISO-PD2: (SP12 wt/wt). Scale bars = 50, 10 and 2 μm respectively. **c.** Representative ICC for DAn health within the tri-culture after 3 weeks, comparing the three different conditions (CTRL: SP11; ISO-PD: ISO-PD1 (SP13 wt/wt); LRRK2-PD: LRRK2-PD1 (SP13)). DAn are stained for TH and mature neurons are stained for MAP2. Scale bar = 25 μm. **d.** Quantification of percentage of TH/MAP2 within the tri-culture after 3 weeks, comparing the three conditions (CTRL: SP11; ISO-PD: ISO-PD1 (SP13 wt/wt) and ISO-PD2 (SP12 wt/wt); LRRK2-PD: LRRK2-PD1 (SP13) and LRRK2-PD2 (SP12)). N=3-5 for ISO-PD1/LRRK2-PD1 cultures and N=3 for ISO-PD2/LRRK2-PD2 cultures. Individual data plotted, along with mean ± SEM. Ordinary one-way ANOVA with Tukey’s multiple comparisons test. **e.** Representative ICC images for DAn (TH+) neurite length in the tri-culture after 3 weeks, comparing the three different conditions (CTRL: SP11; ISO-PD1: SP13 wt/wt; LRRK2-PD1: SP13). **f.** Quantification of DAn neurite length within the tri-culture after 3 weeks, comparing the three conditions (CTRL: SP11; ISO-PD: ISO-PD1 (SP13 wt/wt) and ISO-PD2 (SP12 wt/wt); LRRK2-PD: LRRK2-PD1 (SP13) and LRRK2-PD2 (SP12)). N=2-3 for ISO-PD1/LRRK2-PD1 cultures and N=2 for ISO-PD2/LRRK2-PD2 cultures. At least 35 neurons considered per N. Individual data plotted, along with mean ± SEM. Kruskal-Wallis test with Dunn’s multiple comparisons test. **g.** Representative ICC images for MG (IBA1+) number and morphology within the tri-culture after 3 weeks, comparing the control condition with the LRRK2-PD Astrocytes condition (CTRL: SP11; ISO-PD1: SP13 wt/wt; LRRK2-PD1: SP13). **H.** Quantification of MG (IBA1+) number within the tri-culture after 3 weeks, comparing the control condition with the LRRK2-PD Astrocytes condition (CTRL: SP11; ISO-PD1: SP13 wt/wt; LRRK2-PD1: SP13). N=4. Individual data plotted, along with mean ± SEM. Paired t-test. **i.** Quantification of MG (IBA1+) circularity within the tri-culture after 3 weeks, comparing the control condition with the LRRK2-PD astrocytes condition (CTRL: SP11; ISO-PD: ISO-PD1 (SP13 wt/wt) and ISO-PD2 (SP12 wt/wt); LRRK2-PD: LRRK2-PD1 (SP13) and LRRK2-PD2 (SP12)). N=3 for ISO-PD1/LRRK2-PD1 cultures and N=1 for ISO-PD2/LRRK2-PD2 cultures. At least 100 MG considered per N. Box and whiskers plot with 10-90^th^ percentile. Mann-Whitney test. For all depicted graphs *p<0.05, **p<0.01, ***p<0.001, ****p<0.0001. p-value is specified for values between 0.05 and 0.1. p-values over 0.1 (non-significant) are not shown.

**Table 1.**
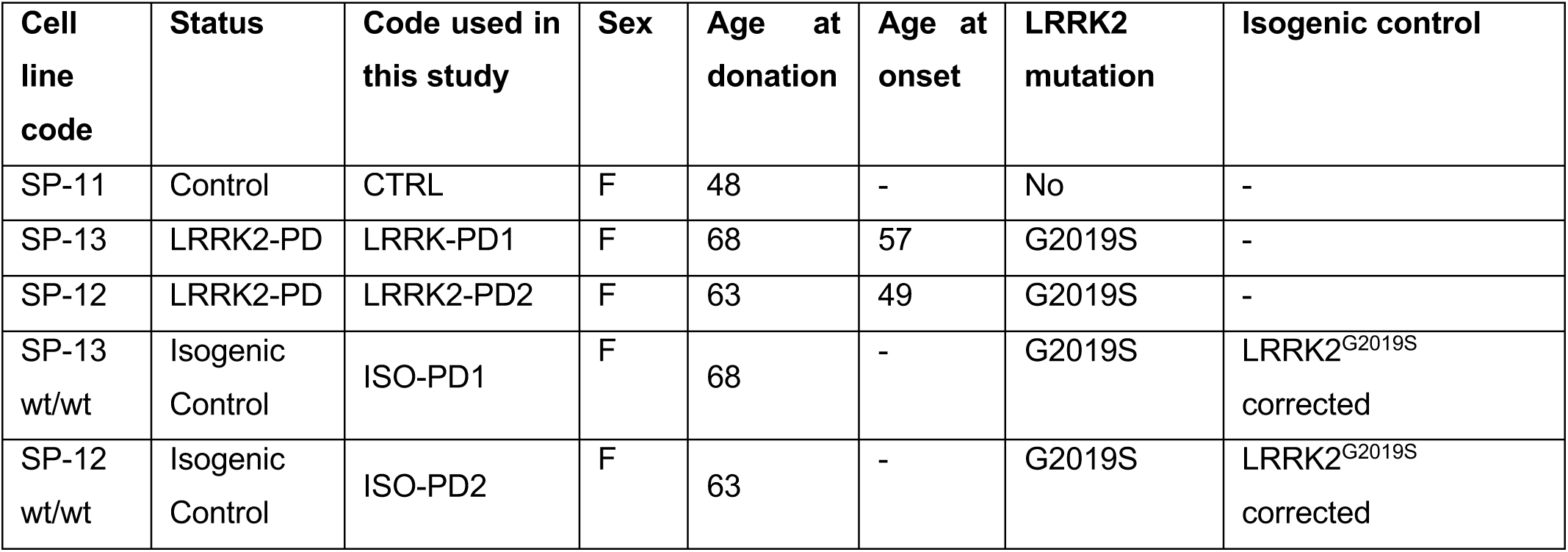
Summary of healthy controls and patients’ lines used in this study.

| Cell line code | Status | Code used in this study | Sex | Age at donation | Age at onset | LRRK2 mutation | Isogenic control |
| --- | --- | --- | --- | --- | --- | --- | --- |
| SP-11 | Control | CTRL | F | 48 | - | No | - |
| SP-13 | LRRK2-PD | LRRK-PD1 | F | 68 | 57 | G2019S | - |
| SP-12 | LRRK2-PD | LRRK2-PD2 | F | 63 | 49 | G2019S | - |
| SP-13 wt/wt | Isogenic Control | ISO-PD1 | F | 68 | - | G2019S | LRRK2<sup>G2019S</sup> corrected |
| SP-12 wt/wt | Isogenic Control | ISO-PD2 | F | 63 | - | G2019S | LRRK2<sup>G2019S</sup> corrected |

To establish the tri-culture system, day 35 DAn were plated onto a monolayer of mature astrocytes, followed by the addition of mature MG one week later. The cultures were then maintained for either one- or three-weeks post-MG addition (**Figure 1A**). We first validated the stability of this system at both timepoints. Under control conditions (CTRL DAn with ISO-PD MG and ISO-PD astrocytes), the three cell types remained uniformly distributed across the culture surface. Astrocytes and neurons formed a dense, interconnected network, while MG were observed resting on top and actively interacting with this glial-neuronal substrate. By week 3, all cell types were in close physical proximity and exhibited clear signs of interaction (**Figure 1B**, **Movie S1**).

To further assess system stability, we quantified the relative proportions of each cell type at both timepoints and compared them to initial plating ratios. Astrocytes, neurons, and MG were plated at a 4:10:5 ratio. These proportions remained remarkably stable throughout the culture period, with observed ratios approximating 5:9:5 at week 1 and 3:9:7 at week 3 (**Figure S1C**). Overall, these data show that our tri-culture system provides a robust and reproducible *in vitro* platform for studying neuro-glial interactions. The system maintains stable cellular proportions and a structurally integrated network over time, enabling the investigation of complex cellular dynamics relevant to PD pathogenesis.

#### Astrocyte-driven microglial activation contributes to dopaminergic neuron loss in a *LRRK2*-PD model

After establishing and characterizing the tri-culture system, we leveraged it to investigate how the *LRRK2*-G2019S mutation in either MG or astrocytes affects DAn health. Specifically, we compared a control condition (ISO-PD MG and ISO-PD astrocytes in contact with CTRL DAn) with a *LRRK2*-PD MG condition (CTRL DAn, *LRRK2*-PD MG, ISO-PD astrocytes) and a *LRRK2*-PD astrocyte condition (CTRL DAn, ISO-PD MG, and *LRRK2*-PD astrocytes). After one week of tri-culture, we did not observe significant differences in DAn number across the three conditions (**Figure S2A-B**). Similarly, DAn neurite length did not differ significantly, indicating that DAn are not visibly affected by the presence of *LRRK2*-PD mutant glial cells at this early timepoint (**Figure S2C**). This lack of a visible DAn phenotype at an early stage is consistent with previous results from our laboratory using DAn-astrocytes co-cultures^16^, and prompted us to assess DAn health after three weeks of tri-culture.

By three weeks, both *LRRK2*-PD mutated glial cells induced a reduction in DAn number, with *LRRK2*-PD astrocytes causing a significantly greater DAn loss (**Figure 1C-D**). Importantly, the number of mature MAP2-positive neurons remained unchanged across the three conditions, demonstrating that *LRRK2*-PD astrocytes specifically induce the degeneration of TH-positive neurons (**Figure S1D**). To further validate these observations, we measured DAn neurite length. Consistent with the DAn cell counts, neurite length was reduced both in the presence of *LRRK2*-PD MG and *LRRK2*-PD astrocytes, with the most pronounced reduction observed when neurons were cultured with the latter (**Figure 1E-F**). These results suggest a major contribution of *LRRK2*-PD astrocytes to DAn degeneration in our tri-culture system. Thus, we hypothesized that this effect may not originate solely from direct astrocyte-to-DAn interaction, but also from astrocyte-mediated changes in the MG phenotype, which could in turn contribute to DAn damage.

To assess whether *LRRK2*-PD astrocytes induce changes in MG phenotype, we compared microglial number and morphology between the control condition and the *LRRK2*-PD astrocyte condition. By the third week of culture, we observed a notable increase in microglial proliferation and a shift toward a rounder morphology in the presence of *LRRK2*-PD astrocytes, a phenotype associated with reactive microglia (**Figure 1G-I**). Consistent with our observations on neuronal changes, the increase in MG proliferation was only detectable at the third week, while no differences were observed at the first timepoint (**Figure S2D**). However, we detected early increases in pro-inflammatory cytokines induced by *LRRK2*-PD astrocytes by the first week of tri-culture (**Figure S2E**), suggesting an early MG activation phase followed by the acquisition of a reactive phenotype at later timepoints. Finally, to validate *LRRK2*-PD astrocyte-induced MG reactivity, we assessed microglial expression of OPN, a well-established disease-associated MG (DAM) marker^28^. OPN expression was significantly increased within MG in the presence of *LRRK2*-PD astrocytes, further supporting our hypothesis (**Figure S2F-G**). In summary, our results demonstrate that *LRRK2*-PD mutated astrocytes are key glial drivers of DAn degeneration and MG reactivity, highlighting their central role in the neurodegenerative process within our *in vitro* model.

#### Single-cell RNA sequencing reveals astrocyte-driven dysregulation of glial crosstalk

After establishing the major role of *LRRK2*-PD astrocytes within our tri-culture system, we employed scRNA-seq to transcriptionally validate our findings and uncover potential therapeutic targets for PD. scRNA-seq was performed on all three conditions at both week 1 and week 3 (**Figure 2A**). Through cell annotation, we identified a total of 17 clusters, including cycling cells (CC_1, CC_2, FP_CC), neural progenitor cells (NPC_1, NPC_2, NPC_3, FP_NPC), neuronal cells (Neuro_1, Neuro_2, Neuro_3, Neuro_4, FP_Neuro), astrocytes (Astro_1, Astro_2), microglia (MG), other glial cells (Other_Glia), and oligodendrocyte progenitor cells (OPC) (**Figure 2B**). Clusters were annotated according to well-established cell-type markers, and clusters pertaining to the floor plate lineage (FP) were identified based on their expression of *OTX2*, a transcription factor known to play a key role in floor plate specification^29–31^ (**Figure 2C**). Importantly, we confirmed an even distribution of cells across clusters in all conditions and timepoints (**Figure S3A**). To visualize DAn, we sub-clustered all neuronal cells and repeated cell annotation (**Figure 2D** and **Figure S3B**). We identified a DAn cluster expressing *TH, PBX1, KCNJ6, LMX1B, and NR4A2*, consistent with previously published hiPSC-derived DAn datasets^31^, as well as a subtype of floor plate progenitor cells (FP_RG) expressing *LMX1A* and *LMO3* (**Figure 2E**).

**Figure 2.**
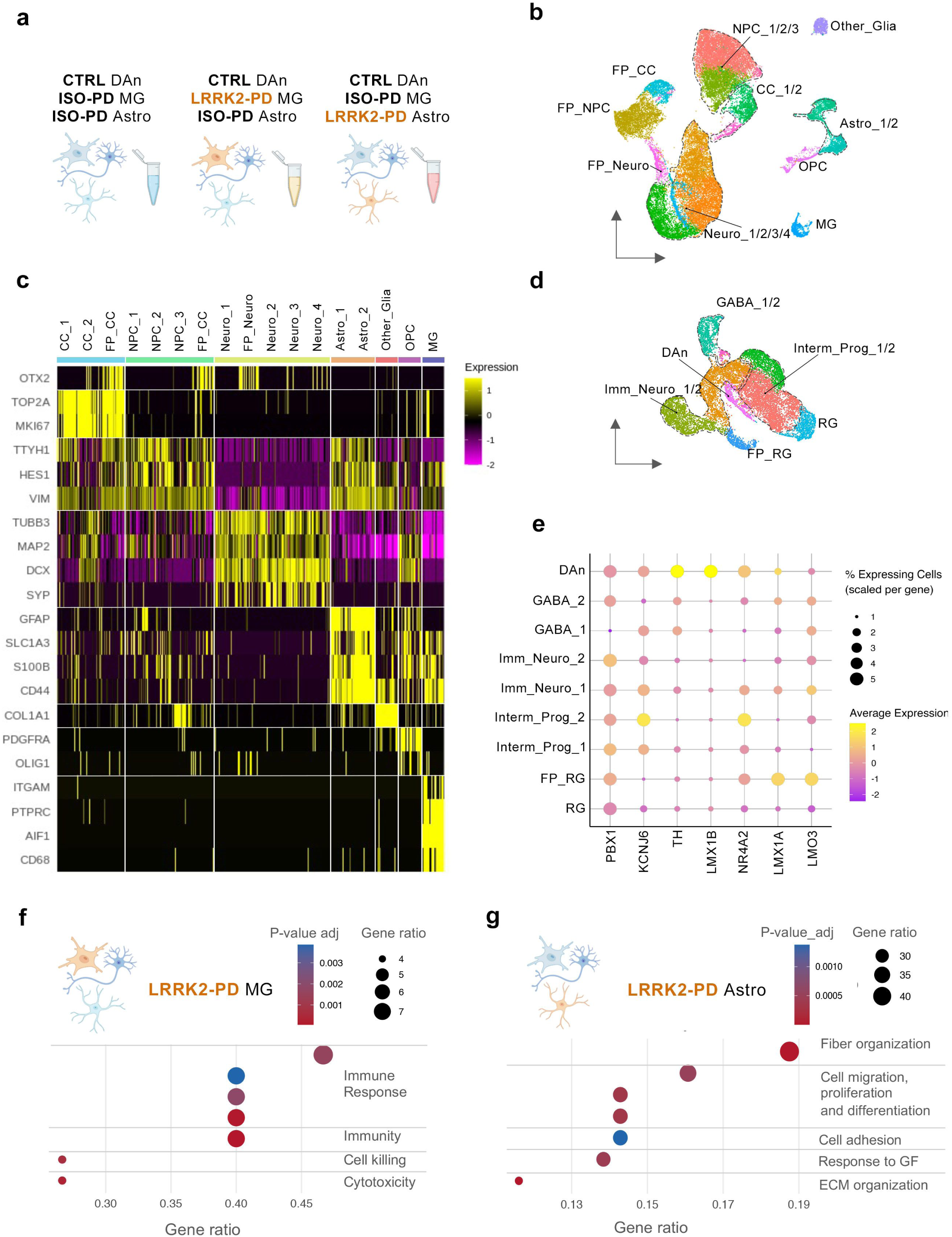
Transcriptional analyses of the tri-culture system. **a.** Schematic representation of the conditions analysed with scRNAseq. **b.** Cluster annotation of the UMAP generated merging the three conditions (CTRL: SP11; ISO-PD (ISO-PD1: SP13 wt/wt); LRRK2-PD (LRRK2-PD1: SP13)) at both timepoints (1 and 3 weeks of tri-culture). **c.** Heatmap displaying the markers used to define each cluster. **d.** Cluster annotation of the UMAP generated subclustering the neuronal cell populations, merging the three conditions (CTRL: SP11; ISO-PD1: SP13 wt/wt; LRRK2-PD1: SP13) at both timepoints (1 and 3 weeks) **e.** Bubbleplot displaying the expression of known Dan markers across the clusters of the neuronal UMAP. **F.** Upregulated biological processes (BP) in LRRK2-PD (LRRK2-PD1: SP13) MG after 1 week of tri-culture, if compared to ISO-PD (ISO-PD1: SP13 wt/wt) MG. **g.** Upregulated BP in LRRK2-PD (LRRK2-PD1: SP13) astrocytes after 1 week of tri-culture, if compared to ISO-PD (ISO-PD1: SP13 wt/wt) astrocytes.

Having confirmed that scRNA-seq captured all cell types of interest, we assessed the impact of *LRRK2*-mutated glial cells on DAn. To do so, we performed enrichment analyses of differentially expressed genes (DEGs) in the DAn cluster upon the presence of either *LRRK2*- PD MG or *LRRK2*-PD astrocytes. Notably, *LRRK2*-PD MG did not produce a significant transcriptional perturbation in CTRL DAn, with only three significant DEGs at both timepoints (**Table S1-2**). In contrast, a substantially greater transcriptional perturbation was observed at both timepoints in DAn co-cultured with *LRRK2*-PD astrocytes (**Table S3-4**). Enrichment analysis of the DEGs in this condition by the latest timepoint revealed dysregulation of pathways related to neuronal and axonal projection, glucose metabolism, and hypoxia response (**Table S5**), highlighting a potential astrocyte-mediated effect on neuronal integrity, metabolism, and ultimately survival.

Following confirmation of the impact of *LRRK2*-PD astrocytes on DAn health at the transcriptional level, we assessed the transcriptional changes in MG and astrocytes carrying the *LRRK2*-PD mutation. *LRRK2*-PD MG exhibited increased biological processes related to the immune response, consistent with a reactive, activated phenotype (**Figure 2F** and **Figure S3C**). This dysregulated transcriptome, including terms related to cytotoxicity, may explain the mild neurodegeneration observed at week 3 when DAn are in contact with mutated MG (**Figure 1C-F**). Conversely, *LRRK2*-PD astrocytes upregulated biological pathways related to extracellular matrix (ECM) and fiber organization, growth, proliferation, and differentiation (**Figure 2G** and **Figure S3D**). This analysis, together with the increased mRNA expression of GFAP in *LRRK2*-PD astrocytes (log_2_foldchange: 2.0907 and p_value adjusted: 6.50E-39 at week 3), suggests that they undergo reactive astrogliosis upon mutation^32^.

To investigate how the *LRRK2*-PD mutation in glial cells alters their crosstalk, we used LIANA^+^, a recently developed tool for inferring cell-to-cell communication^33^, to assess ligand–receptor interactions across the three conditions at the latest timepoint. To visualize if the effects of mutated glia extended beyond their own signalling, we manually compiled and visualized the interaction counts (**Figure 3A**). The number of microglial total interactions increased in the presence of *LRRK2*-PD MG. Similarly, astrocytes carrying the *LRRK2*-PD mutation exhibited the highest number of outgoing interactions. Notably, *LRRK2*-PD astrocytes not only increased their interactions with neurons and MG but also appeared to trigger an elevated number of MG–neuron interactions. These findings suggest that mutated astrocytes play a central role in reshaping the communication network within the system, likely by inducing a reactive state in MG, which in turn enhances their crosstalk with neurons. The top 10 ligand-receptor interactions inferred using LIANA^+^ in *LRRK2*-PD MG or *LRRK2*-PD astrocytes conditions are represented in **Figure S3E-F**.

**Figure 3.**
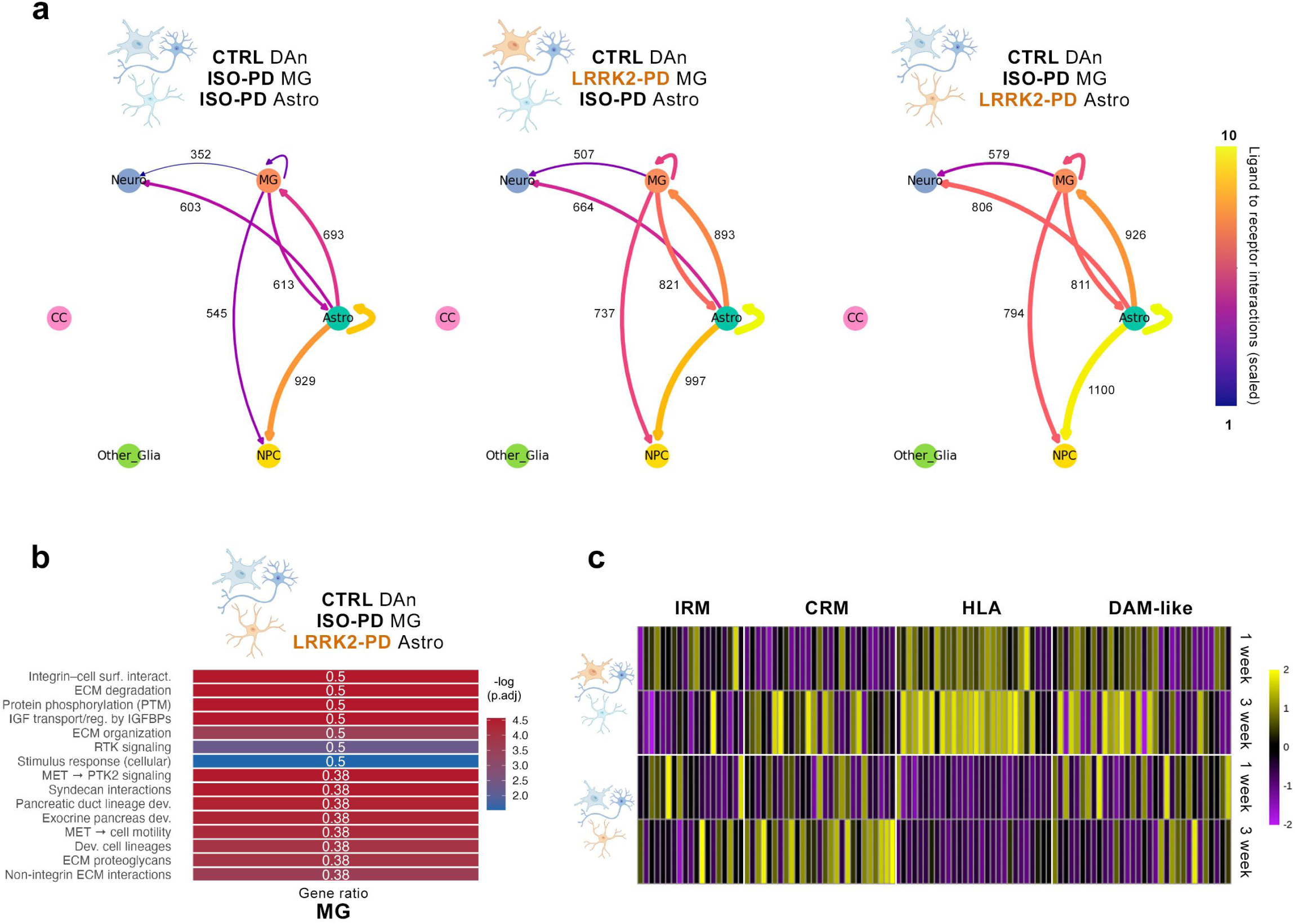
Cellular crosstalk analyses within the tri-culture system. **a.** CirclePlots displaying the number of total ligand-receptors interaction calculated using LIANA+, comparing the three conditions (CTRL: SP11; ISO-PD (ISO-PD1: SP13 wt/wt); LRRK2-PD (LRRK2-PD1: SP13)) after 3 weeks of culture. **b.** Top 15 Reactome upregulated pathways in MG (ISO-PD (ISO-PD1: SP13 wt/wt) upon LRRK2-PD (LRRK2-PD1: SP13) astrocytes presence after 1 week of tri-culture, if compared to control condition (ISO-PD1 MG and ISO-PD1 astrocytes). **C.** Heatmap showing the expression of genes defining IFNy responsive MG (IRM), cytokine response MG (CRM), HLA and DAM-like transcriptional MG profiles^34^ in our dataset, comparing all conditions CTRL: SP11; ISO-PD (ISO-PD1: SP13 wt/wt); LRRK2-PD (LRRK2-PD1: SP13)) at both available timepoints (1 and 3 weeks of tri-culture). Marker genes were taken from a previously published MG atlas^34^. We considered only human genes, ranked as the top 100 markers of that cell state in the original study, and found in at least three datasets.

To further characterize the crosstalk between glial cells upon *LRRK2*-PD mutation, we analysed the DEGs in astrocytes upon *LRRK2*-PD MG presence and in MG when astrocytes are *LRRK2*-PD mutated. *LRRK2*-PD MG did not induce major transcriptional changes in astrocytes, with minimal differences in mRNA levels if compared to the control condition at both timepoints (less than absolute log_2_fold changes of 0.58, **Table S6-7**). In contrast, astrocytes carrying the *LRRK2*-PD mutation markedly reshaped the transcriptional landscape of MG (**Table S8-9**). This response is characterized by the upregulation of Reactome pathways related to microglial reactivity and tissue remodelling, including integrin-mediated cell surface interactions and ECM degradation and organization (**Figure 3B**).

To better identify the phenotype of astrocyte-influenced MG, we compared their transcriptome to that of MG carrying the *LRRK2*-G2019S mutation, using as a reference a recently published MG transcriptional atlas^34^ (**Figure 3C**). While intrinsically mutated MG showed a clear upregulation of genes associated with the HLA-reactive phenotype, isogenic corrected MG in contact with *LRRK2*-PD astrocytes displayed a distinct and time-dependent response. At week 1, their gene expression profile only partially overlapped with interferon gamma–responsive MG (IRM) and DAM-like MG, whereas, by week 3, they more closely resembled inflammatory, cytokine-response MG (CRM). Among the most upregulated genes in this condition was *SPP1* (*OPN*, log_2_foldchange: 2.5143 and p_value adjusted: 1.95E-10 at week 3), which we previously found upregulated at the protein level when MG are in contact with *LRRK2*-PD astrocytes.

In summary, our scRNA-seq data transcriptionally validate the central role of *LRRK2*-mutated astrocytes in driving MG reactivity and remodelling the glial communication network, ultimately contributing to neurodegeneration in our *in vitro* PD model. Moreover, these analyses have identified *SPP1/OPN* as a potential therapeutic target to counteract the observed aberrant astrocytic inflammatory cascade.

#### Ceruloplasmin deficiency in *LRRK2*-PD astrocytes promotes iron dysregulation and oxidative stress

Having established that *LRRK2*-PD astrocytes are key drivers of both microglial activation and DAn degeneration, we next focused on identifying candidate dysregulated molecular targets in this cell population. To this end, we filtered DEGs in *LRRK2*-PD astrocytes based on both statistical significance and effect size, selecting those with an adjusted p-value less than 0.05 and an absolute log₂ fold change greater than 0.58. This yielded 450 DEGs at week 1 and 435 DEGs at week 3 (**Table S10-11**). To prioritize stable and potentially impactful genes, we intersected the two DEG lists, retaining only those consistently dysregulated at both timepoints, resulting in a final set of 209 genes (**Figure 4A**). We then performed pathway enrichment analysis on this gene set, which confirmed that *LRRK2*-PD astrocytes exhibit transcriptional alterations affecting axon guidance and neuronal system development. Moreover, the observed changes were consistent with known consequences of *LRRK2* kinase dysfunction, supporting the pathogenic relevance of the model. Strikingly, among the most significantly enriched terms were those directly or indirectly related to iron homeostasis, including selenoacid metabolism, Diamond-Blackfan anemia, and broader categories of iron dysregulation, suggesting a direct link between the *LRRK2*-PD mutation in astrocytes and disruptions in iron metabolism (**Table S12**).

**Figure 4.**
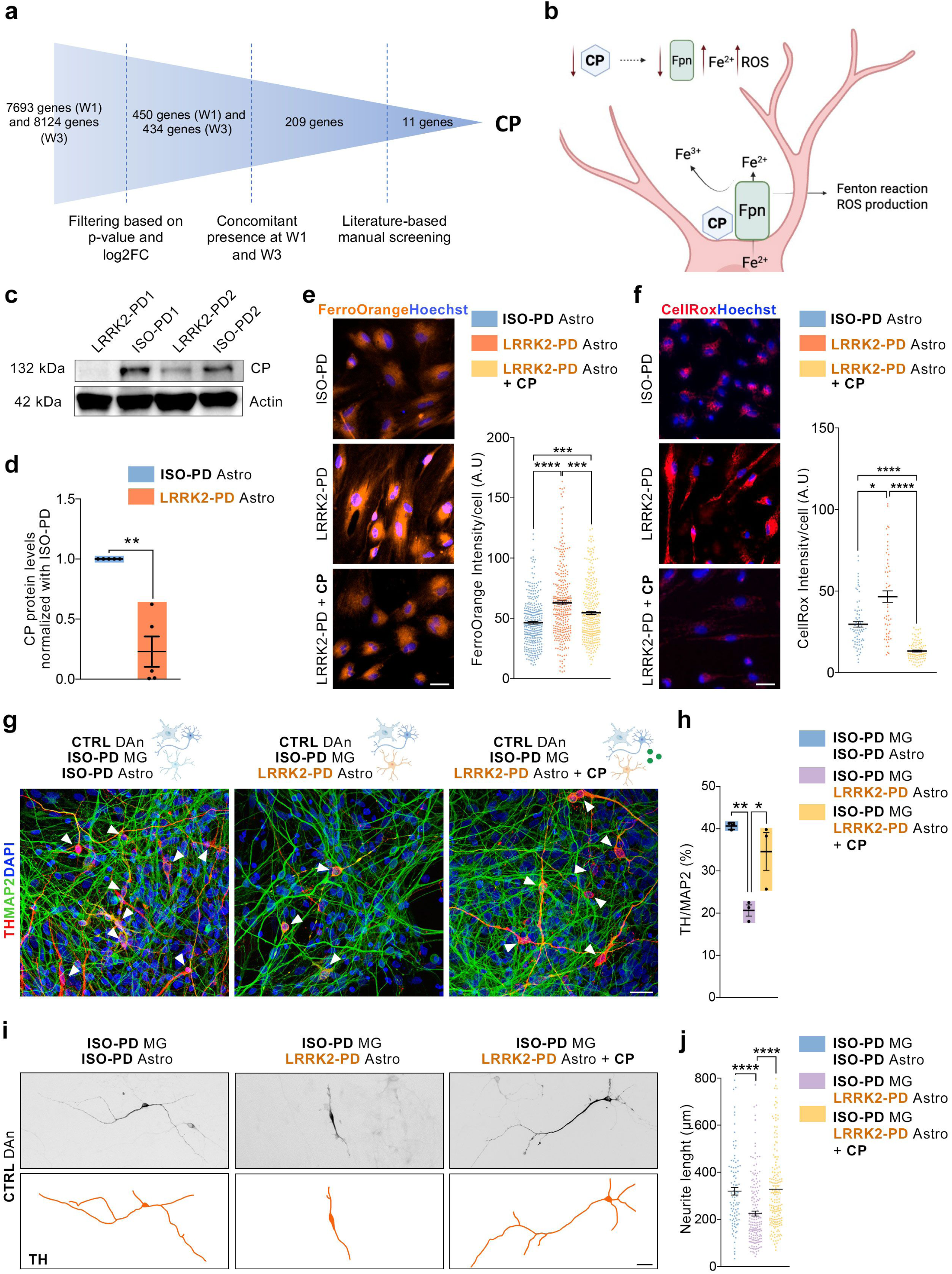
CP downregulation induces iron and ROS accumulation in LRRK2-PD astrocytes, and CP pharmacological treatment prevents Dan from degeneration. **a.** Filtering process followed for the identification of *CP* as a potential molecular target in LRRK2-PD astrocytes. **b.** Schematic representation of the expected consequences of CP downregulation in LRKK2-PD astrocytes. **c.** WB displaying CP levels in the two lines of LRRK2-PD mutated astrocytes (LRRK2-PD1: SP13 and LRRK2-PD2: SP12) if compared to their respective isogenic controls (ISO-PD1: SP13 wt/wt and ISO-PD2: SP12 wt/wt). **d.** Quantification of CP band intensities (normalized on Actin band intensities) comparing LRRK2-PD astrocytes (LRRK2-PD1: SP13 and LRRK2-PD2: SP12) with ISO-PD astrocytes. N=3 for LRRK2-PD1/ISO-PD1 astrocytes and N=2 for LRRK2-PD2/ISO-PD2 astrocytes. Individual data plotted, along with mean ± SEM. One-sample t-test. **e.** Representative images and quantification for intracellular Fe^2+^ using FerroOrange staining comparing ISO-PD astrocytes (ISO-PD1: SP13 wt/wt and ISO-PD2: SP12 wt/wt), LRRK2-PD mutated astrocytes (LRRK2-PD1: SP13 and LRRK2-PD2: SP12) and LRRK2-PD mutated astrocytes treated with CP at 20 μg/mL. Representative images are from LRRK2-PD2 (SP12) and ISO-PD2 (SP12 wt/wt) astrocytes cultures. Scale bar = 20 μm. In the quantification, individual data plotted, along with mean ± SEM. N=3 for LRRK2-PD1/ISO-PD1 and N=3 for LLRK2-PD2/ISO-PD2 astrocytes. At least 35 cells considered per N. Kruskal-Wallis test with Dunn’s multiple comparisons test. **f.** Representative images and quantification for intracellular ROS levels using CellRox staining comparing ISO-PD astrocytes (ISO-PD1: SP13 wt/wt), LRRK2-PD mutated astrocytes (LRRK2-PD1: SP13) and LRRK2-PD mutated astrocytes treated with CP at 20 ug/mL. Scale bar = 40 μm. In the quantification, individual data plotted, along with mean ± SEM. N=2 for LRRK2-PD1/ISO-PD1 astrocytes. At least 25 cells considered per N. Kruskal-Wallis test with Dunn’s multiple comparisons test. **g.** Representative ICC for DAn health within the tri-culture after 3 weeks, comparing the control condition, the one with LRRK2-PD astrocytes, and the one with LRRK2-PD astrocytes treated with CP at 20 μg/mL (CTRL: SP11; ISO-PD: ISO-PD1 (SP13 wt/wt); LRRK2-PD: LRRK2-PD1 (SP13)). DAn are stained for TH and mature neurons are stained for MAP2. Scale bar = 25 μm. **h.** Quantification of percentage of TH/MAP2 within the tri-culture after 3 weeks, comparing the control condition, the one with LRRK2-PD astrocytes, and the one with LRRK2-PD astrocytes treated with CP at 20 μg/mL (CTRL: SP11; ISO-PD: ISO-PD1 (SP13 wt/wt) and ISO-PD2 (SP12 wt/wt); LRRK2-PD: LRRK2-PD1 (SP13) and LRRK2-PD2 (SP12)). N=1 for ISO-PD1/LRRK2-PD1 cultures and N=2 for ISO-PD2/LRRK2-PD2 cultures. Individual data plotted, along with mean ± SEM. Ordinary one-way ANOVA with Tukey’s multiple comparisons test. **i.** Representative ICC images for DAn (TH+) neurite length in the tri-culture after 3 weeks, comparing the control condition, the one with LRRK2-PD astrocytes, and the one with LRRK2-PD astrocytes treated with CP at 20 μg/mL (CTRL: SP11; ISO-PD1: SP13 wt/wt; LRRK2-PD1: SP13). Scale bar = 20 μm. **j.** Quantification of DAn neurite length within the tri-culture after 3 weeks, comparing the control condition, the one with LRRK2-PD astrocytes, and the one with LRRK2-PD astrocytes treated with CP at 20 μg/mL (CTRL: SP11; ISO-PD: ISO-PD1 (SP13 wt/wt) and ISO-PD2 (SP12 wt/wt); LRRK2-PD: LRRK2-PD1 (SP13) and LRRK2-PD2 (SP12)). N=1 for ISO-PD1/LRRK2-PD1 cultures and N=2 for ISO-PD2/LRRK2-PD2 cultures. At least 35 neurons considered per N. Individual data plotted, along with mean ± SEM. Kruskal-Wallis test with Dunn’s multiple comparisons test. For all depicted graphs *p<0.05, **p<0.01, ***p<0.001, ****p<0.0001. p-value is specified for values between 0.05 and 0.1. p-values over 0.1 (non-significant) are not shown.

To further investigate this link, we refined our list, selecting only the genes that have been previously associated with iron and/or ferroptosis. Among the 11 genes selected (**Figure 4A**), ceruloplasmin (*CP*), a copper-dependent iron oxidase, was the most significantly downregulated. Notably, *CP* dysregulation has been previously associated with PD, with decreased protein levels reported in patients’ blood and cerebrospinal fluid (CSF)^35,36^, and astrocytes are the primary source of *CP* synthesis in the brain^37,38^. As *CP* removes Fe^2+^ from the Fenton reaction, acting as an antioxidant, its deficiency should induce an increase in ferrous ion levels coupled with a rise in ROS levels. Moreover, it has been reported that *CP* deficiency induces degradation of ferroportin^39^, which exports Fe^2+^ out of the cell under homeostatic conditions. Given this evidence, we hypothesized that, due to *CP* downregulation, ferrous ions and ROS would accumulate within *LRRK2*-PD astrocytes (**Figure 4B**). To validate these scRNA-seq analyses, we performed qPCRs for iron-related genes in astrocyte monocultures, comparing *LRRK2*-PD with ISO-PD astrocytes. We observed significant downregulation of *CP* and *FTL* and a trend toward downregulation for the other evaluated genes (**Figure S4A**). We then validated *CP* protein-level downregulation by Western blot (WB) in the two *LRRK2*-PD astrocyte lines compared to their isogenic controls (**Figure 4C-D**). To assess the consequences of *CP* deficiency in astrocytes and directly link this deficiency to the observed phenotype, we evaluated Fe^2+^ and ROS levels within astrocytes under basal conditions and upon *CP* pharmacological supplementation. We confirmed an increase in accumulated ferrous ions in *LRRK2*-PD astrocytes, which was partially reversed by *CP* treatment (**Figure 4E**). A similar trend was observed for ROS accumulation, with increased ROS levels in *LRRK2*-PD astrocytes that were significantly decreased by *CP* supplementation (**Figure 4F**). Together, these data confirm a strong downregulation of *CP* in *LRRK2*-PD- mutated astrocytes and implicate a critical role for *CP* in Fe^2+^ and ROS accumulation, thereby contributing to oxidative stress and possibly neurodegeneration in PD.

#### Ceruloplasmin supplementation reverses astrocyte-mediated neurodegeneration and microglial activation

Having established the consequences of *CP* downregulation in astrocytes and demonstrating that *CP* supplementation rescues these changes in astrocyte monocultures, we hypothesized that the accumulation of Fe^2+^ and ROS within astrocytes may be a key driver of the DAn degeneration observed in the tri-culture. To address this hypothesis, we treated the tri-culture system with *CP*, starting at week 2 and continuing until the end of week 3. Strikingly, *CP* supplementation prevented DAn death in the tri-culture, partially restoring DAn numbers to levels comparable to the control condition (**Figure 4G-H**). Importantly, the treatment also prevented neurite degeneration (**Figure 4I-J**).

We also evaluated MG health upon treatment, and, intriguingly, *CP* supplementation reduced MG proliferation and microglial *OPN* intensity, reverting them to levels like those observed in the control condition (**Figure S4B-D**). These results strongly suggest that the aberrant phenotype triggered by *CP* downregulation in *LRRK2*-PD astrocytes is also driving MG activation.

Therefore, we propose a mechanism in which *LRRK2*-PD astrocytes downregulate *CP*, leading to the accumulation of ferrous ions and ROS both within the astrocytes themselves and, potentially, in the surrounding microenvironment. These changes result in DAn degeneration and the adoption of a reactive, inflammatory phenotype in MG. Importantly, both the DAn and MG phenotypes are rescued upon *CP* treatment, demonstrating a clear and direct link between these phenotypes and *CP* downregulation (**Figure S4E**). In summary, our results highlight a strong implication of astrocyte-derived Fe^2+^ and ROS in *LRRK2*-PD pathogenesis and strongly support *CP* as a potential therapeutic target for this disease.

## Discussion

PD, the second most common neurodegenerative disorder, represents a growing global health challenge as populations age^1,40^. Despite advances in understanding PD pathogenesis, the lack of disease-modifying therapies underscores the urgent need to identify targetable pathways. While previous studies have examined the non-cell-autonomous effects of *LRRK2*- PD MG or astrocytes on DAn^16,22,24,41^, the interplay between these glial cell types and their collective impact on neurodegeneration has remained largely unstudied. To address this, we established a novel human iPSC-derived tri-culture system integrating astrocytes, MG, and DAn. Unlike existing models^42,43^, our platform employs a lower MG density, enhancing scalability for parallel experiments, and enables selective genetic manipulation of astrocytes and MG to dissect LRRK2-dependent mechanisms with precision.

Our findings identify *LRRK2*-PD astrocytes as primary drivers of both MG activation and DAn degeneration. While earlier studies demonstrated neurotoxic effects of mutated astrocytes on DAn^16,24,41^, we show these cells exert stronger toxicity than mutant MG, a finding we attribute to their dual role: directly damaging DAn and indirectly amplifying neurodegeneration via microglial priming. Although MG are known to induce reactive astrocytes in PD^44–46^, evidence for the reverse (astrocyte-driven MG activation) has been limited^47^. Our work fills this gap, revealing a central role for astrocytes in *LRRK2*-PD pathology.

Single-cell transcriptomics further illuminated this hierarchy, uncovering that *LRRK2*-mutant astrocytes exhibit a reactive signature consistent with prior reports^24^, and induce profound transcriptional changes in DAn, an effect absent with *LRRK2*-mutant MG alone. Strikingly, *LRRK2*-mutant astrocytes also reprogrammed MG toward a reactive, inflammatory state distinct from their intrinsic LRRK2-linked HLA-driven phenotype. This suggests that in patients (where both glial cells carry the *LRRK2*-PD mutation) astrocytes may amplify microglial neurotoxicity, exacerbating disease progression.

Leveraging our model, we identified CP downregulation as a key mechanistic node. While CP deficiency is documented in PD patient SNpc and CSF^35,36,48^, its role has been controversial^49^. Our data reconcile this by demonstrating that astrocytes (the primary source of CP in the brain^38^) reduce CP expression when carrying the *LRRK2*-G2019S mutation, leading to iron dyshomeostasis (Fe^2+^/ROS accumulation). This aligns with CP’s known function in mitigating Fenton chemistry^50^, and with previous studies linking CP loss to parkinsonism in animal models^51^. We propose that iron-mediated toxicity triggers MG activation and DAn death, potentially via ferroptosis, a pathway increasingly implicated in PD^52^, and now under therapeutic exploration^53^. Critically, CP supplementation rescued all disease phenotypes, confirming its causal role and therapeutic potential.

## Limitations of the study

While our tri-culture system offers unprecedented insight into neuro-glial interactions and crosstalk, its prolonged maturation time and inter-experimental variability required rigorous validation. To mitigate these issues, we performed all key analyses using two independent patient-derived iPSC lines and biological triplicates for each experiment, ensuring the robustness of our findings. Moreover, our transcriptomic analyses yielded relatively low cell numbers for certain populations, particularly DAn, which may have obscured subtle MG-driven transcriptional effects, though key findings were functionally validated.

## Acknowledgments

The authors are indebted to the patients with PD who have participated in this study. The authors thank Meritxell Pons-Espinal for helping with astrocyte generation. Research from the authors’ laboratories is supported by the Spanish Ministry of Science and Innovation-MICINN (PID2019-108792GB-I00, PID2020-116339RB-I00, PID2021-123925OB-I00, PID2022-137963OB-I00 and PID2022-139546OB-I00 supported by MCIN/AEI/10.13039/501100011033 and FEDER, and PDC2021-121051-I00 supported by MCIN/AEI/10.13039/501100011033 and by the European Union Next Generation EU/ PRTR); AGAUR (2021-SGR-974), the Marató de TV3 Foundation (202012-32 and 202331-30); CERCA Program / Generalitat de Catalunya); La Caixa Bank Foundation, Spain (CaixaResearch Health 2024, under the agreement LCF/PR/HR24/00788; CaixaImpulse Innovation 2024 (CI24-20499)). J.M.-M. and VB were recipients of FPI pre-doctoral fellowships (PRE2022-104573 and PRE2020-094465, respectively) from the Spanish Ministry of Economy and Competitiveness (MINECO). M.P. work was supported by a Ramón y Cajal contract of the Spanish Ministry of Science, Innovation and Universities (Grant: RYC2018- 024564-I funded by MICIU/AEI /10.13039/501100011033 and by “El FSE invierte en tu futuro”). F. A. work was supported by a predoctoral contract of the Spanish Ministry of Science, Innovation and Universities (Grant: PRE2020-094049 funded by MICIU/AEI /10.13039/501100011033 and by “FSE invierte en tu futuro”). VT received a pre-doctoral La Caixa INPhINIT Incoming Fellowship (code: LCF/BQ/DI21/11860038). AC was recipient of an ICREA “Academia” Award (Generalitat de Catalunya). We acknowledge the Région Île-de-France for funding support toward the acquisition of the spectral flow cytometer SONY ID7000 at the Cytometry Platform of the SFR Necker. Illustrations were created with Biorender (https://www.biorender.com).

## Author contributions

A.C. conceived the study, managed the project progress, and coordinated the experiments and analyses; V.T., J.M.-M., V.B., M.A., I.F.-C., G.S., N. C.-C., Y.R-P., C.W., J.A., performed the experiments and L.M., M.P., F. A., M. D. and I.F. contributed to data analyses; S.P., A.C., A.R., E.T. provided resources. The paper was prepared by V.T., A.R., S.P. and A.C. with feedback from all authors. All authors have read and approved the current version of the manuscript.

## STAR*METHODS

### RESOURCE AVAILABILITY

#### Lead contact

Further information and requests for resources and reagents should be directed and will be fulfilled by the lead contact, Antonella Consiglio.

## Materials availability

Sharing hiPSC lines used in this study requires approval by the Barcelona Stem Cell Bank owing to country and institution ethical regulations.

## Data and code availability

- This paper does not report original code.
- Any additional information required to reanalyze the data reported in this paper is available from the lead contact upon request.

## EXPERIMENTAL MODEL AND SUBJECT DETAILS

### hiPSC cultures

hiPSC lines were previously generated and fully characterized^16,25^. In this study, we employed one control (CTRL) line (SP11), from a healthy individual, and two LRRK2-PD patient lines (SP13 (LRRK2-PD1) and SP12 (LRRK2-PD2)), together with their respective isogenic controls (SP13 wt/wt (ISO-PD1) and SP12 wt/wt (ISO-PD2) (**Table 1**). The generation and use of human iPSCs in this work were approved by the Spanish competent authorities (Commission on Guarantees concerning the Donation and Use of Human Tissues and Cells of the Carlos III National Institute of Health). All procedures adhered to internal and EU guidelines for research involving derivation of pluripotent cell line. Informed consent was obtained from all patients using forms approved by the Ethical Committee on the Use of Human Subjects in Research at Hospital Clinic in Barcelona.

Before being used for following experiments, hiPSC were thawed on Matrigel (Corning) coated plates and in mTesR-1 medium (Stem Cell Technology), supplemented with 10 μM Rho-associated kinase (ROCK) inhibitor (Miltenyi Biotec; Y27632; RI). The day after thawing, media was fully changed and hiPSC were maintained in mTesR-1, which was changed once per day, until they reached appropriate confluency for differentiation.

## METHOD DETAILS

### Astrocyte generation

Astrocytes were generated from hiPSC as previously described^16,54^. Briefly, hiPSC were cultured and differentiated into spherical neural masses (SNMs), were grown in suspension for 28 days with induction medium ((Dulbecco’s Modified Eagle Medium F12 (DMEM/F12) (Invitrogen), 1% N-2 supplement, 0.1% B27 supplement (Invitrogen), 1% nonessential amino acids (NEAA; Corning), 1% penicillin/streptomycin (PenStrep (P/S); PS-B LabClinics), 1% Glutamax (Invitrogen)) supplemented with 20 ng/mL Leukemia Inhibitory Factor (LIF, MilliporeSigma) and 20 ng/mL Epidermal Growth Factor (EGF, R&D Systems), and again for a further 21 days with propagation medium (DMEM/F12, 1% N-2 supplement, 0.1% B27 supplement, 1% NEAA, 1% PenStrep, 1% glutamax) containing 20 ng/mL Fibroblast Growth Factor 2 (FGF-2, PeproTech) and 20 ng/mL EGF. Finally, SNMs were dissociated into a monolayer, plated on matrigel-coated plates, cultured for 14 days in propagation medium and then for another 14 days in Ciliary Neurotrophic Factor (CNTF) medium (Neurobasal (Invitrogen), 1% Glutamax, 1% P/S, 1% NEAA, 0.2% B27 supplement, 10 ng/mL CNTF (Prospec Cyt-272)). Astrocytes were then plated on Thermanox plastic coverslips (Thermo Fisher Scientific) coated with Matrigel in 24-well plates (for tri-culture and monoculture experiments) or in 6-well plates for qPCR or WB experiments.

### Microglia Generation

MG was generated by following an already published protocol^55,56^. Briefly, single hiPSC colonies were passaged in mTeSR-1 media onto fresh Matrigel-coated 6 cm^2^ plastic plate and grown to the desired size (for 2-4 days). At day (D) 0, the protocol started by supplementing mTeSR-1 medium with 80 ng/mL of Bone Morphogenetic Protein (BMP)-4 (PeproTech^®^), until D4. From D5, SP34 medium (StemProTM-34 SFM (Gibco™), with 1% of P/S and 1% of Glutamax) was employed. For two days, SP34 medium was supplemented with 80 ng/ml of Vascular Endothelial Growth Factor (VEGF, PeproTech^®^), 100 ng/ml of Stem Cell Factor (SCF, PeproTech^®^) and 25 ng/ml of FGF-2 (PeproTech^®^). At D7, 10 and 14 the medium was changed with SP34 supplemented with 50 ng/ml of Fms-like tyrosine kinase 3-Ligand (Flt3-L, PeproTech^®^), 50 ng/ml of IL-3 (PeproTech^®^), 50 ng/ml of SCF (PeproTech^®^), 5 ng/ml of Trombopoietin (TPO, PeproTech^®^) and 50 ng/ml of Macrophage Colony Stimulating Factor (M-CSF, PeproTech^®^). Starting from D14, media was changed every 3-4 days with SP34 supplemented with Flt3-L, M-CSF and 25 ng/ml of granulocyte-macrophage colony-stimulating factor (GM-CSF, PeproTech^®^). From D35 onwards, floating MG progenitors were collected from the culture’s supernatant and passed through a 70 μm Filcon™ Syringe-Type nylon mesh (BD Biosciences) to ensure a single-cell suspension and to remove cell clumps.

Cells were counted, centrifuged at 300*g* for 10 minutes and cultured in Roswell Park Memorial Institute (RPMI) 1640 Medium (Gibco™) with 1% of P/S and 1% of Glutamax, supplemented with 50 ng/ml M-CSF and 50 ng/ml IL-34 (PeproTech^®^). Medium was changed with fresh factors every 2-3 days. MG progenitors were deemed as mature MG after 7 days in culture. To plate them into the tri-culture, they were detached using Accutase, counted and plated on top of astrocytes-neuron co-cultures.

### Dopaminergic neuron generation

Ventral midbrain DAn were differentiated by adapting previously published protocols^26,57^. Briefly, at D0 (established as the day where hiPSC cultured on Matrigel-coated plates in mTesR-1 reach 70% of confluence,) mTesR-1 was replaced by NMM media (1:1 DMEM-F12 and Neurobasal, 0.25% Insulin (Sigma Aldrich), 0.5% of Pyruvate Sodium (Gibco™– Termo Fisher Scientific), P/S, N2, NEEA and Glutamax, 2% B27-VitA (Invitrogen), 1:1000 of b- Mercaptoethanol (Gibco™ – Termo Fisher Scientific)). From D11 to D15, NMM media was substituted with N2B27 media (Neurobasal, 1% B27-VitA, 0.5% N2, and 1% of P/S, NEAA and Glutamax). Finally, from D16 onwards, the media used was the terminal differentiation one (Neurobasal, 2% B27-vitA, 1% of P/S and Glutamax). Supplemented factors included: 10 mM SB (from D0 to D7, PeproTech^®^), 500 nM LDN (from D0 to D12, PeproTech^®^), 200ng/mL to 1µM Sonic Hedgehog (SHH) (from D0 to D9, PeproTech^®^), 1.5µM to 3µM CHIR (from D3 to D12, PeproTech^®^), 100ng/ml FGF8b (from D9 to D16, PeproTech^®^). From D16 onwards Brain-Derived Neurotrophic Factor (BDNF) (20 ng/mL, PeproTech^®^), Glial cell line-Derived Neurotrophic Factor (GDNF (20 ng/mL, PeproTech^®^), TGFβ3 (1 ng/mL, PeproTech^®^), DAPT (10 mM, PeproTech^®^), ascorbic acid (0,2 mM, PeproTech^®^) and AMPc (0,1 nM, PeproTech^®^) were used. Media was changed every other day. At D22 of differentiation, neurons were passaged on poli-D Lysine:Laminine coated 24 well plate with glass coverslip for immunocytochemistry staining or on coated 6 well-plates for following experiments. D35 mature DAn were detached using Accutase, counted and plated on top of astrocytes for tri-culture experiments.

### Triple culture set-up

To generate the tri-cultures, 20.000 mature astrocytes were first seeded at day 0 onto 24 well plates Thermanox plastic coverslips coated with matrigel and cultured using CNTF medium for 7 days. At day 7, 50.000 DAn were plated on top of astrocytes. The co-culture was maintained using co-culture media (Neurobasal, 1% Glutamax, 1% P/S, 1% NEAA, 2% B27- VitA supplement). At day 14, 25.000 MG were added, forming the tri-culture, which was maintained in tri-culture media (1:1 Neurobasal:RPMI), supplemented with 50ng/mL IL-34 and 50 ng/ml M-CSF. The media was then changed once every 2-3 days, depending on media consumption. The endpoint for analyses were at 1 and 3 weeks post MG addition. Unless specified differently in figure legend, analyses were performed in tri-cultures coming from two different patients’ lines, and pooled in the quantifications to minimize line-dependent variability.

### CP treatment

CP was supplemented at a final concentration of 20 μg/mL to the cell culture of interest. Supplementation was performed together with media change and twice a week. For astrocytes monocultures, CP was supplemented from D21 to D35 after astrocytes plating. In tri-cultures, CP was supplemented from day 10 up to the third week after MG addition.

### Triple culture dissociation

1 and 3 weeks LRRK2-PD1/ISO-PD1 tri-cultures were dissociated using a papain-based tissue dissociation kit (Worthington LK003150). Cells were disaggregated using a P1000 and filtered using a 40mm cell strainer to reach a single cell suspension. After centrifugation, cells were resuspended in their own media (tri-culture media) and kept in ice until encapsulation. Counting and viability assessment were performed using Trypan blue staining.

### Data processing, alignment and gene quantification

FASTQ files were processed using Cell Ranger (10x Genomic Cell Ranger version 7.0.0). First, a genome index for the hg38 version of the human genome from the 10x Genomics support site was downloaded. Then, *cellranger count* function was used to generate single cell count matrices for each of the samples. A total of 35.385 cells were obtained.

### Quality control and filtering

Single cell count matrices of each sample were imported using the R package Seurat v4.2.0^58^ and merged into a unique Seurat object. Then, Seurat was used to filter low quality cells. First, the percentage of reads associated with mitochondrial genes for each cell was calculated, as high content of these types of RNAs has been previously associated with low quality/damaged cells. Then, cells with less than 1000 Unique Molecular Identifiers (UMIs) and that had a gene count higher than the 95% of all the cells were discarded. Moreover, cells that had a percentage of mitochondrial genes above 10% were filtered out. In addition, a linear model was fitted to describe the relationship between the log number of UMIs and the log number of genes detected per cell. All cells with a residual smaller than −0.5 were discarded. As a result of this second filtering, we started our analyses with a total of 34.524 cells. The final cell count for each sample can be found in **Supplementary Table 13**.

### Data integration, dimensionality reduction and clustering

The Seurat function *SCTransform*^59^ was used to regress out the effect of the UMI counts, gene counts and the percentage of mitochondrial UMIs, and to scale and normalize the data. Additionally, *SCTransform* identified 2.500 highly variable genes for each of the samples. To correct for batch effects, the samples were integrated based on astrocyte condition and timepoint using Canonical Correlation Analysis (CCA), with the function *SelectIntegrationFeatures* to find the integration features to be used. A principal component analysis (PCA) was then performed on the unified Seurat Object. By using the *Elbowplot* function, the eigenvalues corresponding to each individual component were plotted. After manual inspection, the first 11 principal components (PCs) were selected for performing downstream analyses. The clustering analysis was performed by using the function *FindClusters,* with the Louvain algorithm parameter for the modularity optimization with a resolution of 0.4 applied on the KNN tree graph. As a result of the clustering analysis, 17 clusters were obtained.

### Classification of cells based on cell-type specific marker genes

To define the identity of each of the cell clusters, cluster-specific marker genes were generated by performing differential gene expression analysis on the integrated Seurat object. The function *FindAllMarkers* from Seurat, with parameters *min.pct* set to 0.25 and *logfc.threshold* set to 0.25, was employed to identify marker genes. Cluster identification and annotation was done using a set of manually curated genes and the publicly online cell-type marker databases PanglaoDB^60^ and CellMarker website^61^.

### Subclustering of neuronal cell types

To get a characterization of specific neuronal cell types, the clusters corresponding to immature neurons and neuronal cell populations were selected. On these clusters, the normalization using *SCTransform* function was repeated to identify 2.000 variable genes. After the dimensional reduction process, 10 PCs were used for dimensionality reduction and clustering. The cells were clustered by using the function *FindClusters* using a resolution of 0.2, and the process resulted in a total of 9 identified clusters. For the cell type characterization, we identified the markers of each cluster using the *FindAllMarkers* function, and defined cluster identity basing ourself on a set of manually curated genes.

### Identification of differentially expressed genes between conditions

To evaluate the changes in gene expression among different conditions, we performed a differential gene expression analysis for each cluster using the Seurat function *FindMarkers*. Genes with an absolute log_2_fold change >= 0.58, and an adjusted p-value < 0.05 were considered significantly differentially expressed, after having performed a MAST statistical test.

### Enrichment analysis

To identify enriched gene sets associated to a described pathway process among the differentially expressed genes (DEG), we performed a pathway enrichment analysis of the up and down regulated DEG (considering only the ones with adjusted p-value < 0.05) using EnrichR^62^. For generating figures related to up-regulated BP or Reactome pathways in a given condition, we used the *EnrichGo* (pvalueCutoff = 0,01 and qvalueCutoff = 0,05) and the *enrichPathway* (pvalueCutoff = 0,05).

### Cell-to-cell communication analyses

To evaluate cell-to-cell communication we first grouped the different clusters in Neurons (including Neuro_1, Neuro_2, Neuro_3, Neuro_4, and FP_Neuro), NPC (including NPC_1, NPC_2, NPC_3, and FP_NPC), CC (including CC_1, CC_2, and FP_CC), astrocytes (including Astro_1 and Astro_2), MG, and Other_glia (including Other_glia and OPC). We then employed the LIANA+ tool^33^. Analyses were performed with *rank aggregate* function, with default parameters. In circle plot, interaction counts were plotted. Values were normalised on a colour scale with the function: *def scale_raw_counts(count, min_count, max_count, min_width=1, max_width=10): return min_width + (count - min_count) / (max_count - min_count + 1e-5) * (max_width - min_width)*, where min_count and max_count where the minimum and maximum absolute value, respectively.

### Gene expression analyses by qPCR

RNA was extracted using a RNeasy® Mini kit (Qiagen) according to manufacturer’s instructions and quantified with a NanoDrop spectrophotometer (Thermo Fisher Scientific). Reverse transcription into cDNA was performed with the Superscript III First-Strand Synthesis System (InvitrogenTM), and a final concentration of cDNA of 2 ng/μL was obtained. Gene expression was measured by real time quantitative Reverse Transcription Polymerase Chain Reaction (qRT-PCR). 8 ng of cDNA were loaded in every well of a MicroAmpTM Optical 384- well Reaction Plate (Applied BiosystemsTM) with 10 nM of forward and reverse primers (Life Technologies; **Supplementary Table 14**) and PowerUpTM SYBRTM Green Master Mix (Applied BiosystemsTM). Plate was run in a 7900HT Fast Real-Time PCR System with 384- well Block Module (Applied BiosystemsTM) following a standard cycling mode. Cycling values were normalized to the housekeeping gene β-Actin and the relative fold gene expression of samples was calculated using the 2-ΔΔCt method.

### Immunofluorescence

To perform immunofluorescence staining (IF), cells plated in 24-well plates were first fixed using 4% of Paraformaldehyde (PFA; EMS). Briefly, half medium was substituted with PFA and left for 5 minutes at room temperature (RT). This procedure was repeated three times (during the last time PFA was left for 10 minutes at RT). To wash out PFA, three rinses substituting half PFA with Dulbecco’s phosphate-buffered saline (DPBS) were performed.

After fixation, samples were rinsed three times in 1X Tris-Buffered Saline (TBS) with 0.1% of Triton before being blocked at RT for 2 hours with Blocking Solution (TBS with 0.01% of Triton and 3% of Donkey Serum). After that, samples were incubated with primary antibodies at 4°C for 48 hours (**Supplementary Table 15**). Samples were rinsed three times in 1X TBS with 0.1% of Triton and blocked with Blocking solution at RT for 1 hour. Incubation for 2 hours with secondary antibodies at RT followed, whit samples being protected from light. All secondary antibodies employed were from the Alexa Fluor Series (Jackson ImmunoResearch Europe) and were used at a concentration of 1:250. Finally, samples were rinsed in TBS 1X and nuclei were counterstained with 0.5 µg/mL of 4’,6-Diamidino-2-phenylindole (DAPI; Abcam) at RT for 10 minutes. Samples were mounted within a glass coverslip (Corning^®^) with Polyvinyl Alcohol- 1,4-Diazabicyclo-Octane (PVA-DABCO; Sigma-Aldrich). Mounted slides were dried at RT for 30 minutes protected from light and stored at 4°C before analysis.

### ROS staining

ROS were measured using CellRox^TM^ Green reactive (ThermoFisher). Briefly, 5 uM of CellRox was added to the live cultures, which were then incubated at 37 degrees for 30 minutes. The dye was washed with three DPBS rinses, and cells were counterstained with Hoechst to be able to visualize nuclei. ROS were quantified in live cells immediately after dye staining.

### Intracellular ferrous ion staining

Intracellular ferrous ion was measured using FerroOrange (Cell Signalling Technology) and following manufacturer instructions. Briefly, cells were incubated with 1 µmol/L of FerroOrange at 37 degrees for 30 minutes and iron levels were quantified in live cells immediately after. Hoechst dye was added in order to stain nuclei.

### Image acquisition and analysis

Images were acquired using Carl Zeiss Axio Imager M2 with an ApoTome (Zeiss Microscopy) microscope using a Cascade CCD camera (Photometrics). For confocal images, a Carl Zeiss LSM880 confocal microscope with an SPE confocal system was employed (Zeiss Microscopy). Neurite length analysis was performed using the Simple Neurite Tracer Plugin for ImageJ^63^. Microglia morphological analyses were performed using Microglia Morphometry Plugin for ImageJ^64^. To measure ROS, the fluorescence intensity of the CellRox dye was measured for every cell, subtracting the background intensity. To measure ferrous iron concentration, cells dyed with FerroOrange were imaged and the dye intensity was calculated per each cell, subtracting the background of the image. OPN analyses were performed by measuring OPN levels per each cell. For image rendering, Photoshop^®^ (Adobe) and IMARIS (Bitplane copyright) were employed.

### Immunoblot

Cells were solubilized in RIPA buffer (150mM NaCl, 50mM Tris pH=8, 1% NP-40, 0.5% NaDoc, 0.1% SDS) containing proteinase and phosphatase inhibitors (Roche), and protein concentration was determined by Bradford assay. All the fractions were separated by SDS-PAGE and immunoblotting was performed using anti- ceruloplasmin (1:380, Abcam ab48614) and anti-Actin (1:400 Sigma A5228) as primary antibodies followed by mouse and rabbit HRP-linked whole Abs (1:2000, Amersham ECL NA931 and NA934 respectively).

### Cytokine analysis

Cytokine analysis has been performed using the LEGENDplex^TM^ Human Essential Immune Response Panel (BioLegend GmbH, Germany) following the manufacturer instructions. Samples were read using a Sony ID7000^TM^ spectral cell analyzer (Sony Biotechnology, USA). For each target cytokine, more than 500 beads were recorded per sample. Data were analyzed using the LEGENDplex^TM^ Biolegend software.

### Statistical analysis

Statistical analyses were performed using GraphPad Prism version 7.0a for MacOSX, (GraphPad Software, Boston, Massachusetts USA, www.graphpad.com). Outlier values were determined using ROUT test with Q set to 1%. Data normality was assessed with Shapiro-Wilk test. For normally distributed data, pairwise comparisons were done using two-tailed t-test. For data comprising more than two groups, we used one-way ANOVA followed by a post-hoc test, as specified in figure legends. For data departing from normality, we used the Mann-

Whitney test when two groups were compared, or the Kruskal-Wallis followed by a post-hoc test as specified in figure legends when comparing more than two groups.

