## Supplementary Material for "Astrocytic Ceruloplasmin Deficiency Triggers Iron Toxicity and Neurodegeneration in a LRRK2 Parkinson’s Tri-Culture Model"

**Word Count**

| | |
|---|---|
| Summary word count | 154 words |
| Title length | 108 characters |
| Total word count (excluding STAR methods and references) | 5620 words |
| Main Figures | 4 |
| Main Tables | 1 |
| Supplemental Figures | 4 |
| Supplemental Tables | 15 |
| References | 64 |

### Abstract

Astrocytes and microglia carrying the LRRK2-G2019S mutation contribute to non-cell-autonomous dopaminergic neuron (DAn) degeneration in Parkinson's disease (PD), but the mechanisms underlying their interplay remain unclear. Here, we developed a novel induced pluripotent stem cell (iPSC)-derived tri-culture system comprising healthy DAn and either LRRK2-mutant or isogenic control iPSC-derived astrocytes and microglia. Using integrated functional assays and transcriptomic profiling, we found that mutant astrocytes adopt a hyperreactive state, driving microglial activation and subsequent DAn degeneration. Mechanistically, we identified a selective downregulation of ceruloplasmin (CP), a copper-dependent ferroxidase, in mutant astrocytes, leading to disrupted iron homeostasis with accumulation of $Fe^{2+}$ and ROS. This iron dysregulation mediated both microglial reactivity and neurodegeneration. Notably, pharmacological restoration of CP reestablished iron homeostasis, reduced microglial activation, and protected DAn from degeneration. Our findings uncover a novel astrocyte-microglia-neuron axis driving PD pathogenesis and showcase the power of our unique stem cell tri-culture platform for dissecting disease mechanisms and discovering therapeutic targets.

### Keywords

Parkinson's disease, LRRK2-G2019S, astrocytes, microglia, dopaminergic neurons, ceruloplasmin, iron homeostasis, iPSC, tri-culture model, neurodegeneration

**Figure S1**

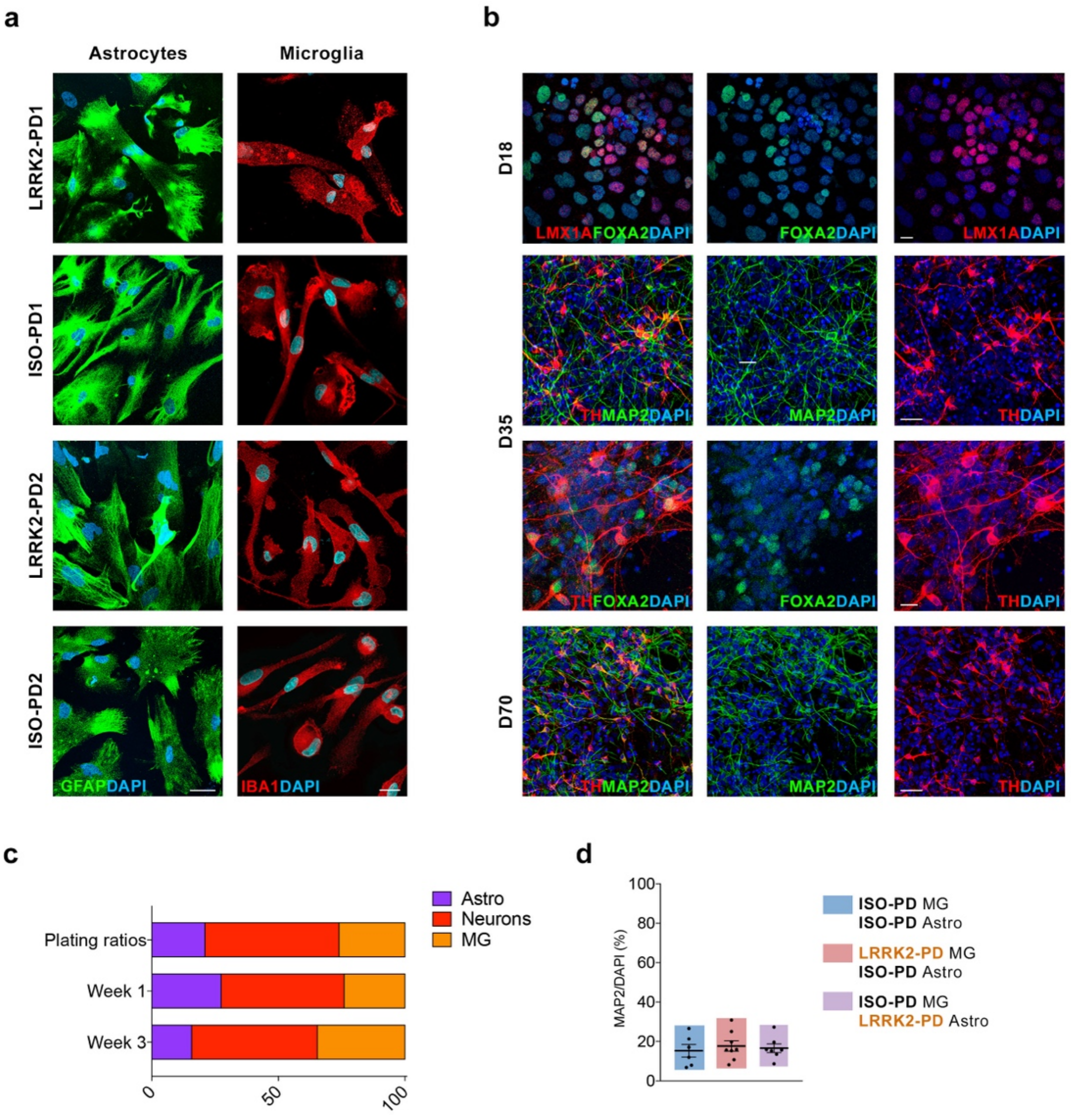


**Supplementary Figure 1: Characterisation of astrocytes, MG, and DAn**
**a.** Representative ICC images for mature astrocytes and mature MG (marked respectively with GFAP and IBA1) generated from the four available hiPSCs lines (LRRK2-PD1: SP13, ISO-PD1: SP13 wt/wt, LRRK2-PD2: SP12, ISO-PD2: SP12 wt/wt). Scale bar = 20 and 10 µm respectively. **b.** Representative ICC images of: floor plate committed neural progenitor cells (NPC, D18), staining positive for LMX1A and FOXA2. Scale bar = 5 µm; mature (D35) DAn, staining positive for TH and MAP2 (Scale bar = 50 µm), and TH and FOXA2 (Scale bar = 1 µm); D70 DAn, staining positive for TH and MAP2. Scale bar = 50 µm. **c.** Comparison between plating ratios in the triple culture and astrocytes:Neurons:MG ratios after 1 or 3 weeks of triple culture. Ratios are taken from an ISO-PD1 (SP13 wt/wt) MG, ISO-PD1 (SP13 wt/wt) a-strocytes and CTRL (SP11) DAn culture (N=1). Similar results are observed in all the generated triple cultures. **d.** Quantification of the MAP2/DAPI percentage in the triple culture after 3 weeks, comparing the three conditions (CTRL: SP11; ISO-PD: ISO-PD1 (SP13 wt/wt) and ISO-PD2 (SP12 wt/wt); LRRK2-PD: LRRK2-PD1 (SP13) and LRRK2-PD2 (SP12)). N=3-5 for ISO-PD1/LRRK2-PD1 cultures and N=3 for ISO-PD2/LRRK2-PD2 cultures. Individual data plotted, along with mean ± SEM. p-values over 0.1 (non-significant) are not shown.

**Figure S2**

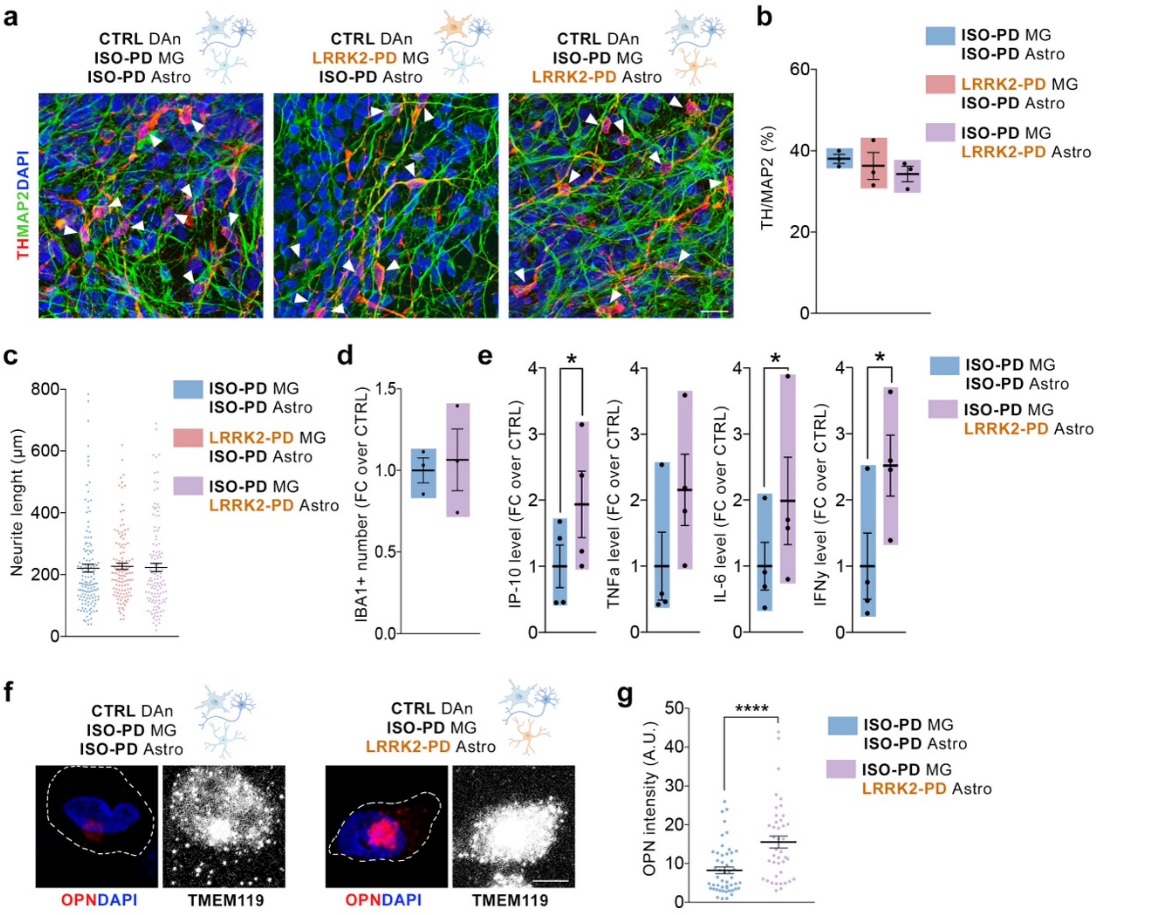


**Supplementary Figure 2: DAn phenotype at 1 week of triple culture and activated microglial phenotype upon contact with LRRK2-PD astrocytes**

**a.** Representative ICC for DAn health within the triple culture after 1 week, comparing the three different conditions (CTRL: SP11; ISO-PD2: SP12 wt/wt; LRRK2-PD2: SP12). DAn are stained for TH and mature neurons are stained for MAP2. Scale bar = 25 µm **b.** Quantification of percentage of TH/MAP2 within the triple culture after 1 week, comparing the three conditions (CTRL: SP11; ISO-PD: ISO-PD1 (SP13 wt/wt) and ISO-PD2 (SP12 wt/wt); LRRK2-PD: LRRK2-PD1 (SP13) and LRRK2-PD2 (SP12)). N=2 for ISO-PD1/LRRK2-PD1 cultures and N=1 for ISO-PD2/LRRK2-PD2 cultures. Individual data plotted, along with mean ± SEM. **c.** Quantification of DAn neurite length within the triple culture after 1 week, comparing the three conditions (CTRL: SP11; ISO-PD: ISO-PD1 (SP13 wt/wt); LRRK2-PD: LRRK2-PD1 (SP13)). N=2-3. At least 35 neurons considered per N. Individual data plotted, along with mean ± SEM. **d.** Quantification of MG (IBA1+) number within the triple culture after 1 week, comparing the control condition with the LRRK2-PD astrocytes condition (CTRL: SP11; ISO-PD: ISO-PD1 (SP13 wt/wt) and ISO-PD2 (SP12 wt/wt); LRRK2-PD: LRRK2-PD1 (SP13) and LRRK2-PD2 (SP12)). N=2 for ISO-PD1/LRRK2-PD1 cultures and N=1 for ISO-PD2/LRRK2-PD2 cultures. Individual data plotted, along with mean ± SEM. **e.** Quantification of pro-inflammatory cytokines levels within media after 1 week of triple culture, comparing the control condition with the LRRK2-PD astrocytes condition (CTRL: SP11; ISO-PD: ISO-PD1 (SP13 wt/wt) and ISO-PD2 (SP12 wt/wt); LRRK2-PD: LRRK2-PD1 (SP13) and LRRK2-PD2 (SP12)). N=2 for ISO-PD1/LRRK2-PD1 cultures and N=2 for ISO-PD2/LRRK2-PD2 cultures. Individual data plotted, along with mean ± SEM. Paired t-test for IP-10, IL-6 and IFNy; Wilcoxon test for TNFa. **f.** Representative ICC images for OPN intensity per microglial cell after 3 weeks of triple culture, comparing the control condition with the LRRK2-PD astrocytes condition (CTRL: SP11; ISO-PD: ISO-PD1 (SP13 wt/wt); LRRK2-PD: LRRK2-PD1 (SP13)). MG is recognized by positivity for TMEM119 staining. Scale bar = 5 µm **g.** Quantification of OPN intensity per microglial cell after 3 weeks of triple culture, comparing the control condition with the LRRK2-PD astrocytes condition (CTRL: SP11; ISO-PD: ISO-PD1 (SP13 wt/wt); LRRK2-PD: LRRK2-PD1 (SP13)). N=3. Individual data plotted, along with mean ± SEM. Mann-Whitney test. *$p<0.05$, ****$p<0.0001$. p-values over 0.1 (non-significant) are not shown.

**Figure S3**

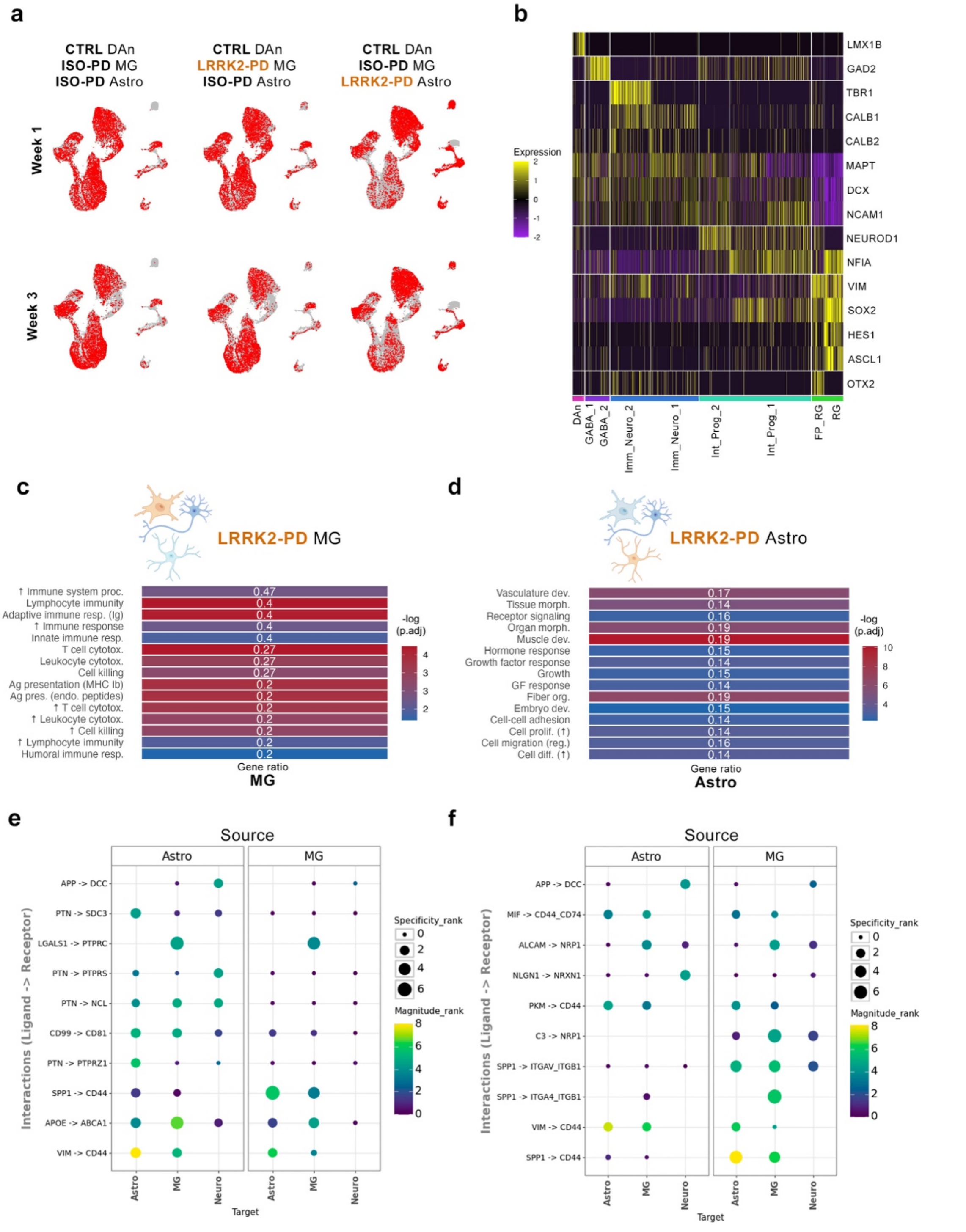


**Supplementary Figure 3: Study of LRRK2-PD mutation impact on glial cells and their crosstalk trough scRNAseq**

**a.** Distribution of cells across all sample in the transcriptomic analyses. **b.** Heatmap plot showing the genes used for cluster annotation of the neuronal sub-clustering. **c.** Top 15 upregulated biological processes (BP) in LRRK2-PD (LRRK2-PD1: SP13) MG after 1 week of triple culture, if compared to ISO-PD (ISO-PD1: SP13 wt/wt) MG. **d.** Top 15 upregulated BP in LRRK2-PD (LRRK2-PD1: SP13) astrocytes after 1 week of triple culture, if compared to ISO-PD (ISO-PD1: SP13 wt/wt) astrocytes. **e.** Bubbleplot displaying the top 10 ligand to receptor interactions coming from astrocytes or MG as a source, and having astrocytes, MG or neurons as a target in the LRRK2-PD MG condition (LRRK2-PD1 (SP13) MG, ISO-PD1 (SP13 wt/wt) astrocytes and CTRL (SP11) DAn. **f.** Bubbleplot displaying the top 10 ligand to receptor interactions coming from astrocytes or MG as a source, and having astrocytes, MG or neurons as a target in the LRRK2-PD astrocyte condition (ISO-PD1 (SP13 wt/wt) MG, LRRK2-PD1 (SP13) astrocytes and CTRL (SP11) DAn.

**Figure S4**

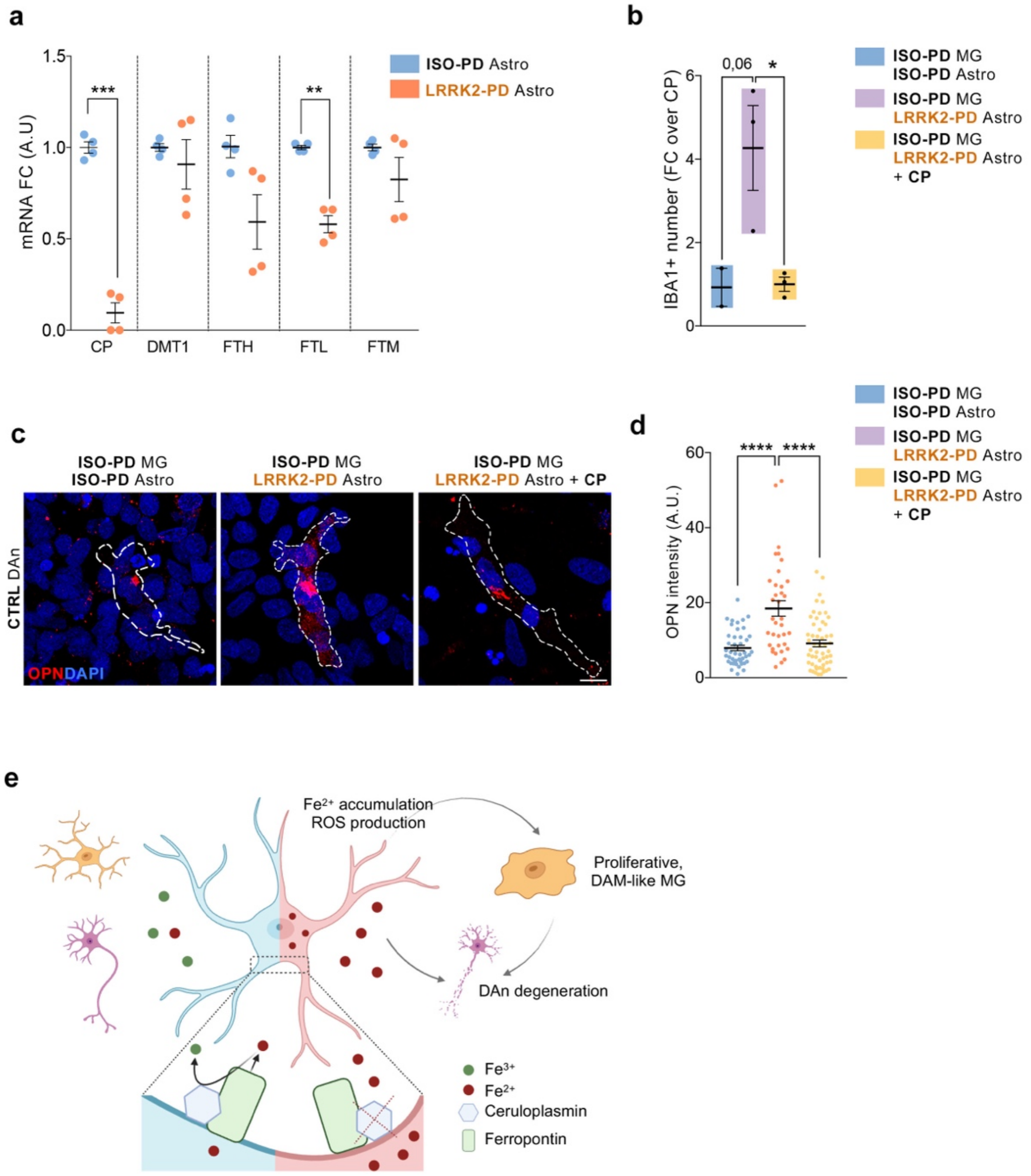


**Supplementary Figure 4: CP supplementation restore LRRK2-PD astrocytes-induced MG reactive phenotype**

**a.** qPCR of iron related genes in astrocytes monocultures, comparing LRRK2-PD (LRRK2-PD1: SP13) with ISO-PD (ISO-PD1: SP13 wt/wt) astrocytes. N=2 with 2 technical replicates per N. Individual data plotted, along with mean ± SEM. One sample t-test. **b.** Quantification of MG (IBA1+) number within the triple culture after 3 weeks, comparing the control condition, the one with LRRK2-PD astrocytes, and the one with LRRK2-PD astrocytes treated with CP at 20 µg/mL (CTRL: SP11; ISO-PD: ISO-PD1 (SP13 wt/wt); LRRK2-PD: LRRK2-PD1 (SP13)). N=3 Individual data plotted, along with mean ± SEM. Ordinary one-way ANOVA with Tukey's multiple comparisons test. **c.** Representative ICC images for OPN intensity per microglial cell after 3 weeks of triple culture, comparing the control condition, the one with LRRK2-PD astrocytes, and the one with LRRK2-PD astrocytes treated with CP at 20 µg/mL (CTRL: SP11; ISO-PD: ISO-PD1 (SP13 wt/wt); LRRK2-PD: LRRK2-PD1 (SP13)). **d.** Quantification of OPN intensity per microglial cell after 3 weeks of triple culture, comparing the control condition, the one with LRRK2-PD astrocytes, and the one with LRRK2-PD astrocytes treated with CP at 20 µg/mL (CTRL: SP11; ISO-PD: ISO-PD1 (SP13 wt/wt); LRRK2-PD: LRRK2-PD1 (SP13)). N=2. At least 10 MG considered per N. Kruskal-Wallis test with Dunn's multiple comparisons test. For all depicted graphs *p<0.05, **p<0.01, ***p<0.001, ****p<0.0001. p-value is specified for values between 0,05 and 0,1. p-values over 0.1 (non-significant) are not shown. **e.** Proposed mechanisms for astrocytes-induced DAn degeneration during LRRK2-PD (red astrocyte) if compared to a healthy condition (blue astrocyte).

**Movie S1:** IMARIS reconstruction of a control triple culture (TH+ DAn (CTRL: SP11), GFP+ MG (ISO-PD2: SP12 wt/wt), and GFAP+ astrocytes (ISO-PD2: (SP12 wt/wt)) showing how the three cell types are in close contact with each other after 3 weeks from MG plating.
